# Using user-centered design to better understand challenges faced during genetic analyses by novice genomic researchers

**DOI:** 10.64898/2026.02.06.704411

**Authors:** Harsh Vijaykumar Patel, Gail P. Jarvik, Taryn Hall, David Veenstra, Serena Jinchen Xie, David Crosslin

## Abstract

The lack of user-centered design principles in the current landscape of commonly-used bioinformatics software tools poses challenges for novice genomics researchers (NGRs) entering the genomics ecosystem. Comparing the usability of one analysis software to that of another is a non-trivial task and requires evaluation criteria that incorporates perspectives from both existing literature and a diverse, underrepresented user base of NGRs. To better characterize these barriers, we utilized a two-pronged approach consisting of a literature review of existing bioinformatics tools and semi-structured interviews of the needs of NGRs. From both knowledge sources, the key attributes that resulted in poor adoption and sustained use of most bioinformatics tools included poor documentation, lack of readily-accessible informational content, challenges with installation and dependency coordination, and inconsistent error messages/progress indicators. Combining the findings from the literature review and the insights gained by interviewing the NGRs, an evaluation rubric was created that can be utilized to grade existing and future bioinformatics tools. This rubric acts as a summary of key components needed for software tools to cater to the diverse needs of both NGRs and experienced users. Due to the rapidly evolving nature of genomics research, it becomes increasingly important to critically evaluate existing tools and develop new ones that will help build a strong foundation for future exploration.

## Background and Significance

Bioinformatics is a rapidly growing field that involves the application of statistical methods and computational algorithms to analyze and interpret biological data. With the increasing availability of genomics data, the development of bioinformatics tools has become essential for researchers to efficiently analyze and interpret the vast amounts of biological data generated. To rival this rapid growth of genomic data, the amount of bioinformatics tools being developed and published has increased dramatically [1]. While most of those tools are developed in siloed laboratories, large scale analysis platforms have also risen in usage and popularity, such as National Human Genome Research Institute’s (NHGRI) Analysis Visualization and Informatics Lab-space (AnVIL)/Terra and the All of Us platforms [2-3]. Most of these tools, however, suffer widespread adoption challenges due to inherent usability barriers from a lack of user-centered design during developed and testing phases.

The International Organization for Standardization defines usability as “the extent to which a product can be used by specified users to achieve specific goals with effectiveness, efficiency and satisfaction in a specified context of use”9. While related fields of study such as global health and healthcare education for medical professionals have enlisted the aid of professional design and innovation firms such as IDEO and Dalberg to tackle challenges that arise with proper usability,[4-5] usability analysis in bioinformatics tools development is scarce and poses many barriers to proper development and sustained adoption [6, 7]. Tools and software packages improperly created without usability considerations in mind suffer from lack of adoption or sustained use [8, 9]. Furthermore, the challenges and distribution of software tools in academia, such as assuming technical background of end users, academic journals and conferences being the primary source of distribution of tools, and the dissemination of incomplete tools, pose great barriers for long-term tool stability.

To address some of the challenges listed above, several bioinformatics tools have been developed specifically with usability and end-user needs in mind. Examples include GenomIUm, where expert bioinformaticians reported reduction of complexity during knowledge extraction and easier analysis workflows, and the All of Us Cohort Selection Pipeline, which allows for streamlined case-control cohort selection [3, 10]. Larger research institutes such as the National Human Genome Research Institute (NHGRI) and EMBL-European Bioinformatics Institute (EMBL-EBI) have also begun to push for more deliberate user-friendly tool development and dissemination citing “the user is the future” [8, 11].

Although there has recently been a push towards developing more user-centered bioinformatics tools, novice genomics researchers (NGR) are overlooked despite being keystone users of these tools. NGRs are individuals who are interested in conducting genetic experiments but lack the quantitative background or experience with existing bioinformatics techniques/tools or do not have access to extensive technological resources, such as local high performance computational clusters. As a core mission of the NHGRI is to train a diverse and talented workforce for the future of genomics in healthcare, NGRs pose as a key user base for the future of bioinformatics tools adoption and sustained usage [11-13]. Machado et al.’s case study on teaching and learning in bioinformatics reinforces the importance of information gathering needs of novice researchers and the hurdles currently faced in the forms of poorly designed bioinformatics tools and databases, lack of proper documentation and support, and vague or missing error handling [14]. UCD principles can be utilized to address these needs of NGRs and produce a workforce capable of interacting with, manipulating, analyzing, and sharing biological data more effectively and efficiently [15].

Albeit limited, some work has begun to address NGR needs. Murphy et al. studied users who regularly navigated complex databases and designed a visual interface to ease the burden for novice users trying to find research patient cohorts [16]. Britto et al. conducted scenario based usability analysis of a pediatric patient portal to study the challenges faced by novice users and reported major barriers of medical language complexity and difficulties in navigation [17]. We even see barriers of information overload and lack of proper tool documentation faced by non-specialists who try utilizing popular ontology creation tools to capture a relatively simple biological system [18, 19]. The challenges faced by novice users in the above studies are not limited to just the contexts of those investigations but are universal across bioinformatics as novice researchers continue to be overlooked.

To fill the gap of usability in bioinformatics tools and to query the needs of NGRs, we employed a mixed-methods study design consisting of two primary parts: a literature review and a needs assessment of target users. First, a retrospective analysis of existing bioinformatics analysis tools utilized by researchers to perform genetics experiments was conducted to better understand the current landscape of available analysis instruments. Thereafter, a prospective analysis using semi-structured interviews was conducted to assess the needs of novice genomics researchers from different sites. We hypothesized the existing literature and the initial needs of NGRs will primarily revolve around thorough documentation and tutorials, minimalistic user-interfaces, and low barriers of installation and deployment of bioinformatics tools.

## Methods

The literature review was conducted using the guidelines set forth by Harris et al [20]. Given the scarcity of user-centered design usage when developing and evaluating bioinformatics tools,[8] specifically tools focused on novice genomics researchers, the scope of the literature review was intentionally left broad in order to best capture as much literary evidence possible for the final analysis and evaluation rubric development.

The primary scope of the literature review was to survey the existing landscape of bioinformatics tools. Specifically, the review centered around the following research question: “What are the available bioinformatics analysis tools commonly utilized to conduct genetic analyses?”.

**Table 1** shows the search strings utilized to identify relevant literature in the surveyed databases:

**Table 1.** Search criteria utilized to find relevant literature from different databases. Except for Pubmed, all other databases were manually filtered for the publication date (post 2003) and English language requirements [1].

| Database | Search String |
| --- | --- |
| Pubmed | ((("bioinformatic") OR ("biomedical informatic") OR ("genomic") OR ("genetic") OR ("informatic") OR ("gene"))) AND ((("software") OR ("tool") OR ("platform") OR ("analysis") OR ("user-friendly") OR ("UCD"))) AND ("2003"[Date - Publication] : "3000"[Date - Publication]) AND (English[Language])) |
| Web of Science | ((("bioinformatic") OR ("biomedical informatic") OR ("genomic") OR ("genetic") OR ("informatic") OR ("gene"))) AND ((("software") OR ("tool") OR ("platform") OR ("analysis") OR ("user-friendly") OR ("UCD"))) |
| Medline | ((("bioinformatic") OR ("biomedical informatic") OR ("genomic") OR ("genetic") OR ("informatic") OR ("gene"))) AND ((("software") OR ("tool") OR ("platform") OR ("analysis") OR ("user-friendly") OR ("UCD"))) |
| Bioinformatics | Individual Searches: "genomic", "gene", "user-friendly", "analysis", "GWAS" |
| Software DB |  |
| Google Scholar | ((("bioinformatic") OR ("biomedical informatic") OR ("genomic") OR ("genetic") OR ("informatic") OR ("gene")) AND (("software") OR ("tool") OR ("platform") OR ("analysis") OR ("user-friendly") OR ("UCD"))) |

After executing each search, abstracts were screened using exclusion criteria to remove irrelevant articles: lacking mention of informatics tools or genetic/genomic data, being a conference abstract, dissertation/thesis, not peer-reviewed, not in English, published before 2004, or missing example results/analysis. Remaining articles were then reviewed in full and excluded if they showed minimal discussion of genomics software, lacked tool URLs, omitted documentation or analysis types, or failed to provide enough detail to identify potential users.

Prior to recruiting participants, an Institutional Review Board (IRB) application for this study was submitted to the University of Washington’s Human Subjects Division on August 31st, 2022. This application was approved on September 7th, 2022 with exempt status. After obtaining approval from the IRB, a target sample of 12 study participants were recruited for interviews. This sample size falls in line with the estimate of thematic saturation typically being reached between 10-20 interviews in studies involving qualitative health research [21-22]. The inclusion criteria for study participants were as follows:

- Graduate student currently belonging to a lab at the University of Washington and Tulane University
- Prior experience with bioinformatics analysis methodologies
- Prior experience with informatics tools and software
- Less than 3 years of experience with genomic analyses

The primary investigator (HP) created preliminary questions for a 30-minute semi-structured interview, which were reviewed by a qualitative expert at UW. Based on feedback, an initial guide was developed and tested in two pilot interviews. A revised, more comprehensive version was then created to address missing concepts and reassessed after two additional interviews. This final guide was used for the remainder of the study.

Each interview was scheduled for 45 minutes, which included time for introductions, background on the purpose of the study, informed consent, and the interview itself. Consent was acquired verbally utilizing a consent guide approved by the IRB. A semi-structured interview methodology was utilized when conducting the interviews. DeJonckheere et al. have shown that semi-structured interviews appear to be a good balance between rigor and flexibility and allow for effective extraction of qualitative themes [23].

After each interview, participants were asked to complete a short demographic questionnaire, approved by the IRB. These audio recordings were stored in a secure UW Zoom-cloud account. The output files (a .MP4a file) were then exported to an encrypted device and then subsequently uploaded to a password-protected automatic transcription service called HappyScribe [24]. After transcription, all recordings were permanently deleted from the transcription service and the transcripts were downloaded onto the same encrypted device for post-processing. Additionally, once the transcripts were collected, all audio files were permanently deleted from the UW Zoom-cloud account. Finally, the transcripts were uploaded to a local ATLAS.ti project for qualitative analysis [25].

In order to analyze the transcripts for emergent qualitative themes, template analysis was conducted [26]. Introduced by Crabtree et al., template analysis is a form of thematic analysis that utilizes hierarchical coding derived from a codebook based on an existing template in addition to allowing for new codes that may appear from the transcripts. In the context of this study, the template that was utilized as the foundational basis for codebook development was the interview guide itself. As the interview guide was slightly altered throughout the course of data collection, the corresponding analysis codebook also went through a series of updates and changes. **Table 2** below shows the finalized codebook that was derived from template analysis and utilized to tag relevant information from each of the transcripts:

**Table 2.** Codebook developed throughout template analysis and subsequently utilized to code interview transcripts.

| Code | Description |
| --- | --- |
| First experiences | describes their first experiences when they got into bioinformatics/genomics research |
| Resources used when getting into field | describes what kinds of resources (textbooks, tutorials, courses) they used to help them get into genomics research and conduct experiments |
| Specific software tool(s) | describes any specific tool or software package used to conduct their experiments/workflows |
| Emotional reaction | describes any emotional reaction to utilizing specific tool (i.e. was satisfied, frustrated because of _____) |
| Cognitive reaction | describes any cognitive reaction to utilizing specific tool (i.e. was not able to understand, understood usage and setup easily) |
| Overall experience | describes any overall experience details that were not captured by the other codes |
| Benefits of specific tool | describes specific benefits in utilizing analysis tool (features, usability, etc.) |
| Barriers of specific tool | describes specific barriers in utilizing analysis tool (shortcomings, usability, etc.) |
| Impact on upkeep/cost | describes any costs on data upkeep or analysis computational costs that were incurred as a result of the tool |
| Overall impact | describes any overall impact details that were not captured by the other codes |
| Suggested improvements | describes what improvements would they like to see in the specific tool or more generally to bioinformatics tools being developed in the future |
| Challenges/concerns with cloud genetics | describes what challenges/concerns exist in the future of performing genomics analysis on the cloud |

To arrive at the finalized codebook above, the primary investigator and an independent coder (JC) worked together to first reach high inter-coder agreement. The metric utilized to assess inter-coder reliability was the Krippendorff’s Cu /cu-alpha coefficient [27-28]. After initial independent coding of a few transcripts and comparing codes, a Krippendorff’s Cu /cu-alpha coefficient of 0.919 was achieved, corresponding to high inter-coder reliability. Once this reliability was achieved, the primary investigator and secondary coder split up the remaining transcripts and coded independently.

To synthesize the knowledge gained from the literature review and needs assessment, an evaluation rubric was developed. Evaluation, within the context of professional assessment, is defined as the systematic determination of the quality or value of something [29]. A good evaluation requires knowledge about the evaluand (what you are evaluating), its background, and its context [30]. For this step, the evaluand was the effectiveness and usability of existing bioinformatics tools within the context of NGRs [31]. Utilizing the methodology for evaluating bioinformatics tools and training practices from Xie et al. and Tracentenberg et al., an initial set of evaluation criteria were created to begin rubric creation [32-33]. These criteria were then adjusted to incorporate the needs of NGRs and the strengths and weaknesses of bioinformatics tools gathered from the literature review. The primary investigator then utilized Markiewicz et al.’s primer on evaluation framework development to create a scoring metric for each criteria [34].

## Results

After searching through the aforementioned databases, a total of 385 potential studies were identified. Of these studies, 196 studies were removed due to being duplicates and 133 studies were removed based on title and abstract screening. Of the 56 remaining full-text studies, 22 were removed due to either a lack of genomic tool availability, lack of documentation, or lack of information regarding the scope and target audience of the tool. A total of 34 studies remained and were utilized in the literature review.

Of the genomics analysis tools from the 34 studies, 32 (94%) had documentation/tutorial content provided, 17 (50%) had a graphical user interface, 8 (24%) were cloud-compatible, 8 mentioned some UCD considerations (24%), and only 7 (21%) studies mentioned a target audience of novice users.

**Table 3** displays demographic information about the participants. In summary, most participants (N = 12) were female (58%), Asian (50%), non-Hispanic/Latino (92%), had less than 2 years of experience working with genetic data (75%), ranged in age from 21 to 48 (mean = 27) years old, and had varied education and familiarity with genetics.

**Table 3.** Participant Demographics from Needs Assessment.

|  |  | N |
| --- | --- | --- |
| Age | Mean | 27 (SD 7) years |
|  | Median | 26 (range 21 – 48) years |
| Sex | Male | 5 (42%) |
|  | Female | 7 (58%) |
| Race | White | 4 (33%) |
|  | African-American | - |
|  | Asian | 6 (50%) |
|  | Multi-racial | 1 (8.5%) |
|  | Other | 1 (8.5%) |
| Ethnicity | Hispanic or Latino | 1 (8%) |
|  | Non Hispanic or Latino | 11 (92%) |
| Education Level | Bachelor's Degree | 6 (50%) |
|  | Master's Degree | 5 (42%) |
|  | Doctor of Medicine | 1 (8%) |
| Familiarity with genetics (out of 5) | Median | 3 (range 1 - 4) |
| Years of experience | Less than 1 year | 4 (33%) |
|  | 1 – 2 years | 5 (42%) |
|  | More than 2 years | 3 (25%) |

Participants’ emotional reactions to first utilizing bioinformatics tools varied when asked “How did you feel when you were working with your analysis tool?”. Some participants reported feelings of satisfaction and excitement:

> *“Well, I was super excited about it, actually. This was actually my first time that I got to look at real data and trying to figure out how to use it and make it useful*.*” (P12)*
>
> *“Definitely felt a sense of like satisfaction that I had kind of worked through and gotten things [SVABA] to work*.*” (P8)*

However, the primary emotions shared by the majority of the participants were frustration, confusion, and being overwhelmed:

> *“Honestly, that was terrible for me* … *a starter, the documentation and the reason you run through the pipeline [PLINK] is it’s really hard, like, for a starter to figure out every component in that. So that’s going to cause a lot of headache*.*” (P2)*
>
> *“Definitely overwhelmed at the beginning, I think, because mostly kind of a lack of domain expertise*.*” (P8)*

The emotional reactions above were mirrored in the participants’ cognitive reaction and understanding when asked “What concerns or questions did you have about utilizing (genomics software) when you first started using it?”.

Most participants reported a lack of domain knowledge and technical knowledge needed to properly utilize their analysis tool:

> *“So, I think that it [Bioconductor GWAS] feels a lot of them feel like a black box, essentially, where at least in my experience, you don’t know what’s going on. You don’t know if it’s working. You put the arguments in and you pray, and then when it finishes, you’ll see if it worked or not*.*” (P6)*
>
> *“It definitely caused a lot of frustrations because of their documentation and some of those error messages. It’s not really clear. When you also Google that [the errors] there are a bunch of other people googling the same things you could find and the answers are actually various*.*” (P2)*

There were a few participants who reported sufficient initial understanding of how their tool worked and increased understanding after following tutorials or working with their tool over the span of the project period:

> *“It [AllOfUs] didn’t really have anything that I thought that they were missing. I thought they were pretty clear on how to use everything and they’re very informative. And then they also add a more detailed help support system in terms of like, if you did get lost in trying to understand how to use one of their specific tasks that they had*.*” (P5)*

When asked “Thinking back, what do you see as positives about utilizing_________(their analysis software)?”, most participants reported they appreciated the provided tutorials and documentation, how efficient and robust their tool was (even when the dataset was very large), and built-in privacy and security precautions:

> *“I think the biggest plus for me is that it has a very robust documentation. It still has some holes in it, I think, but the least of the software tools that I was using. Sometimes it has some gaps in it. At least it’s pretty simple to go from zero to being able to use it*.*” (P6)*

When asked “Thinking back, what do you see as negatives about utilizing_________(their analysis software)?”, most participants reported trouble during initial setup, lack of thorough error messages, lack of user-friendly interfaces, and challenges sharing results outputted to colleagues or other pipelines:

> *“I think often times some tools are set up in a way where it’s not immediately evident what you need to do. It was frustrating because you would submit something and you wouldn’t know for a while whether or not it worked. And so then you get the error or whatever a few hours later and then you have to debug and you feel like you’ve lost a few hours*.*” (P8)*

Despite facing diverse initial barriers to entry, most participants reported they benefited overall from learning how to utilize bioinformatics software. Several suggestions to improve both their experience and the experience of future researchers were provided upon asked the question “Do you have any recommendations for changes or additions to_________(genomics software) that would aid in your research?”. Most participants (9/12) reported the need for better documentation and tutorials (with resources written in layman’s terms), an interactive analysis environment, and clearer error messages:

> *“I expected more documentation on the whole thing. They have some documentation, but they don’t have many examples. So I think they could really improve in that regard*.*” (P3)*
>
> *“I would definitely wanted them to provide more examples of the workflow and how the tutorial would go. Like, if you were doing some genomic analysis and you need to say grab or filter specific regions, what do you do and what do you most of the frequently asked questions or errors*.*” (P2)*

Several participants (4/12) reported need for extra information to be readily available during analysis in some sort of “information hub”:

> *“Honestly, I wish any tool’s methods paper had, like, explain it like I’m five section or something. Because, man, some methods papers are just so convoluted and really difficult to understand, and some are, like, not like that. Also, I think I would have wanted the tool to like point to useful resources, like useful background reading*.*” (P8)*

When asked “As the NHGRI and other genomics research organizations are moving toward cloud computing, what challenges/concerns do you anticipate facing when conducting genomic research in the cloud?”, most participants seemed optimistic about the notion of cloud computing for bioinformatics, but were skeptical about topics such as privacy and security, transition time and training needed, and cost:

> *“From my perspective, I think there are a couple of things I mentioned already. What is the cost structure of it? What do I need to know to set it up correctly from a technical perspective? Those kinds of things, of course, like if you’re using health data, you have to think about HIPAA and you know, proper access control, though I think that that’s pretty well set up or like thought out with those cloud computing things*.*” (P12)*

**Table 4** presents the evaluation rubric, adapted from existing bioinformatics criteria and refined using insights from the literature review and needs assessment. It covers analysis robustness, documentation, data restrictions, accessibility, reproducibility, tool interoperability, and user-centered design.

**Table 4.** Evaluation Rubric for Bioinformatics Tools within the context of NGRs.

| Evaluation Rubric for Bioinformatics Tools | Level 0 | Level 1 | Level 2 | Level 3 |
| --- | --- | --- | --- | --- |
| Criteria | Unacceptable | Marginal | Acceptable | Exceptional |
| Robustness of Genetic Analyses | Only capable of one very specific type of genetic analysis (fixed parameters) and lacks ability to scale | Capable of one or two specific analyses but able to scale to large datasets | Supports many genetic analyses, can scale to large datasets, but lacks clear vision for use of tool | Many genetic analyses supported or able to be supported (users can create/share workflows), clear vision for tool's usage |
| Tutorials/Documentation | Had little/no documentation or tutorials about how to setup or utilize tool to perform analyses | Had documentation/tutorials but was difficult to understand/outdated | Had documentation and tutorials but lacked sample data or references to publications | Clear, updated documentation with examples and sample data along with links to publications that utilized the analysis tool |
| Data Restrictions | Supported only one or two specifically formatted data types | Supported specific formatted data types but did not allow use of new data | Supported specific formatted data types and allowed the upload of new data | Supported robust data types and allowed for the upload of new data |
| Accessibility (Cloud/Free-to-Use/Open-Source) | Inaccessible to the target userbase, behind paywall, proprietary code with little option of how to use (their own platform) | Somewhat accessible to the target audience, not open-sourced, may not be free-to-use | Mostly accessible to the target userbase (some wait if user needed to be validated), some open-source components, and free-to-use | Very accessible tool with both local and cloud-based options, open-source and free-to-use |
| Reproducibility/Interaction with Other Tools | Does not have built in functionality for reproducing experiments or interacting with other software tools | Has built functionality for reproducing experiments but does not interact with other software tools | Has reproducibility functionality built-in and able to interact with other tools but in a very limited range | Excellent built in features for reproducing experiments and is able to interact with other tools with clear guidelines and input/output types |
| UCD Design Considerations (Setup, Usage, Evaluation) | Had little to no usability considerations during the design, development, or evaluation of genetic tool | Had some usability considerations but only during one or two phases of the developmental cycle | Had some usability considerations during most of the development phase but lacked clear interaction with the target user base | Utilized usability considerations during all stages of the development process with clear interaction with target userbase |
| Overall Performance | **Unacceptable** | **Marginal** | **Acceptable** | **Exceptional** |
| Points Required | 0-6 | 6-11 | 12-16 | 16-18 |

## Discussion

Using broad strokes, this investigation captures many of the existing challenges present in the current landscape of bioinformatics software tools, such as missing or incomplete documentation, difficulty in setup/support, challenges in reproducibility, and lack of UCD considerations during design and deployment. However, the richness of the data collected during the literature review and user interviews offer more nuanced perspectives as to why these challenges are relevant, how the challenges relate to one another, and how to potentially overcome them in future bioinformatics tools.

From the results of the literature review, we can easily identify common characteristics shared by most pieces of genomic software that is widely used and highly cited in the bioinformatics space over the past two decades. While almost all tools were open-source and offered some sort of documentation/tutorial content, very few tools noted their target user audience or any component of the UCD framework. On top of that, the tools that did mention UCD principles only did so briefly in the introduction and discussion sections and did not actively incorporate them during the design and deployment of the proposed tool. Studies have shown that poor usability considerations when designing instruments for shared use often leads to failure of sustained use [6]. Furthermore, as only 7 out of the 34 tools evaluated mentioned novice users, most genomics tools currently widely-used and cited fail to align with the NHGRI’s 2020 Strategic Vision, specifically the sections for breaking down barriers that impede genomic research and building/improving a robust foundation for future research [11].

The user interviews conducted in this study offered detailed insight into the challenges routinely faced by NGRs during setup, exploration, experimentation, and sustained use of a variety of different bioinformatics software. Almost all NGRs interviewed reported being overwhelmed and confused during the initial adoption stages and struggled to find adequate resources to help guide their learning process. A key source of this confusion was caused by either lack of domain expertise or insufficient technical knowledge needed to operate the software tool.

Participant 6 sums it up nicely by saying:

> “*So, I think that it [Bioconductor GWAS] feels a lot like a black box, essentially, where at least in my experience, you don’t know what’s going on*. ***You don’t know if it’s working. You put the arguments in and you pray, and then when it finishes, you’ll see if it worked or not***.*” (P6)*

This sentiment of missing information is further supported by the call for more detailed documentation, easy to access tutorial content, and clearer error messages by almost all participants. While solutions regarding missing information for novice users exist already in a variety of other disciplines, bioinformatics software tools, especially those currently introduced to NGRS, are not yet thoroughly polished to support novice users [8, 13].

Additionally, most participants reported challenges in reproducibility and shareability of their findings. While research reproducibility is already seen as a crisis by many investigators, this barrier is especially critical for NGRs. Without proper methods to reproduce experiments, NGRs face increased frustration, delay in software adoption, and an overall deficiency in connecting theoretical concepts to practical applications. Potential solutions to this problem that future software tools can consider include integration of workflow description language (WDL), saving prior experiments in a format that can be rerun, providing guided content as part of the documentation, and having available support beyond documentation (such as support email or forums).

Combining the findings from the literature review and the insights gained by interviewing the NGRs, an evaluation rubric was created that can be utilized to grade existing and future bioinformatics tools. This rubric acts as a summary of key components needed for software tools to cater to the diverse needs of both NGRs and experienced users. Of the key components detailed in Table 4, UCD design considerations with respect to tool setup and detailed supplementary information are the most important considerations that one should incorporate when designing usable software, as reflected by the user interviews. Existing studies that utilize these components efficiently report high throughput of results, better user understanding, and more sustained tool use [16, 18, 19]. As this is an exciting time for genomics research, it becomes increasingly important to critically evaluate existing tools and develop new ones that will help build a strong foundation for future exploration.

Despite the literature review of existing bioinformatics tools covering a wide variety of databases and potential sources of commonly used software, there are several limitations that exist which impact its accuracy. All of the final articles were in English, failing to capture any software tools that were published in non-English journals. Incorporating additional peer-reviewed sources and a different interpretation of the research question may lead to different final studies used.

Although the user interviews gathered rich qualitative data that offered a more nuanced perspective into the challenges faced by NGRs, a relatively small number of participants were interviewed. However, despite this, code groups and thematic domains did not appear to change significantly after 10 participants, indicating theoretical saturation. Furthermore, participants were only selected from the University of Washington and Tulane University and were mostly of self-identified White and Asian descent. Their training and experience may not reflect the views of NGRs from other institutions and backgrounds. For future investigations, it is important to include a more diverse participant pool to capture viewpoints that were not present in this study. Finally, it is important to recognize the nature of qualitative research itself and the impact the primary investigator (H.P.) played during the analysis portion. The final results only represent one possible interpretation of the available data.

## Conclusion

This study underscores a pivotal opportunity for the field of bioinformatics: by embracing user-centered design and addressing the needs of novice genomics researchers, we can build tools that are not only more accessible but also more impactful. Improving usability, documentation, and reproducibility will empower a broader, more diverse community of researchers to contribute to genomic discovery. As genomics continues to evolve, prioritizing usability ensures that innovation is not limited by expertise, but expanded by it, fueling a more collaborative and transformative future.

